# The role of strigolactones in regulation of stomatal conductance and plant-pathogen interactions in *Arabidopsis thaliana*

**DOI:** 10.1101/573873

**Authors:** Maria Kalliola, Liina Jakobson, Pär Davidsson, Ville Pennanen, Cezary Waszczak, Dmitry Yarmolinsky, Olena Zamora, E. Tapio Palva, Tarja Kariola, Hannes Kollist, Mikael Brosché

**Author notes:** Correspondence Mikael Brosché, Viikki Plant Science Centre, Organismal and Evolutionary Biology Research Programme, Faculty of Biological and Environmental Sciences, University of Helsinki, Helsinki, Finland.

## Abstract

Strigolactones are a group of phytohormones that control shoot branching in *Arabidopsis thaliana*. However, in recent years they have been shown to affect many other plant processes. We previously showed that the strigolactone perception mutant *more axillary branches 2 (max2)* has increased susceptibility to plant pathogenic bacteria as a result of more open stomata as well as alterations in hormonal signalling. Here we show that both, strigolactone biosynthesis- (*max3* and *max4*), and perception mutants (*max2* and *dwarf14*) are significantly more sensitive to *Pseudomonas syringae* DC3000. Moreover, in response to *P. syringae* infection, high levels of SA accumulated in *max2* and this mutant was ozone sensitive. To search for the mechanisms that could explain pathogen- and ozone sensitivity we performed gene expression analysis and several different assays that explore the function of guard cells and regulation of guard cell signalling.

Treatments with GR24 (a strigolactone analogue) resulted in very modest changes in defence-related gene expression. In contrast, guard cell function was clearly impaired in *max2* and depending on the assay used, also in *max3, max4* and *d14* mutants. Moreover, stomatal responses to stimuli that cause stomatal closure in wild-type plants (darkness, high CO_2_ and ABA) were analysed in the strigolactone mutants. In darkness both strigolactone biosynthesis and perception mutants showed reduced stomatal closure, whereas the response to high CO_2_ was impaired only in *max2* and *d14*. The response to ABA was not impaired in any of the mutants. To position the role of MAX2 in the guard cell signalling network, *max2* was crossed with mutants defective in ABA biosynthesis (*aba2*), in guard cell ABA signalling (*ost1*) and a scaffold protein required for proper ion channel activity (*ghr1*). The stomatal conductance of double mutants was consistently higher than the corresponding single mutants, suggesting that MAX2 acts in a signalling pathway that functions in parallel to the well characterized guard cell ABA signalling pathway. We propose that the impaired defence responses of *max2* is related to more open stomata that allows increased entry of bacteria or air pollutants like ozone. Furthermore, as MAX2 appears to act in a specific branch of guard cell signalling (related to CO_2_ signalling), this protein could be one of the elusive components that allow guard cells to distinguish between different environmental conditions.

## Introduction

Strigolactones are best known for their role in regulation of shoot branching in *Arabidopsis thaliana* by influencing polar auxin transport (Hayward et al., 2009, Crawford et al., 2010). However, in addition to shoot development, strigolactones affect several other processes such as senescence and responses to salinity and light stress (Shen et al., 2007, Umehara et al., 2008, Gomez-Roldan et al., 2008). Moreover, recently Stes et al., (2015) demonstrated that strigolactones contribute to tolerance to the leafy gall syndrome (caused by *Rhodococcus fascians*), which indicates their role in regulating plant-pathogen interactions. Strigolactones also affect drought tolerance by affecting stomatal conductance (Bu et al., 2014, Ha et al., 2014).

Strigolactones are synthesized mainly in roots and transported to shoots, however the route of transport is not clear (Kohlen et al., 2011; Xie et al., 2015). Both strigolactone synthesis and perception involve *MORE AXILLARY GROWTH (MAX)* proteins that act in a single pathway. MAX1, MAX3 and MAX4 catalyse strigolactone biosynthesis, while the perception require MAX2 and DWARF14 (D14) - the receptor of strigolactone (Al-Babili and Bouwmeester, 2015, Chevalier et al., 2014, Waters et al., 2012). MAX2 is an F-box protein that targets proteins for destruction as part of the ubiquitin proteasome system (Lechner et al., 2006, Stirnberg et al., 2002). Strigolactones are proposed to be hydrolyzed by D14 resulting in a conformational change in the D14 protein, which facilitates interaction of D14 and SCF^MAX2^ (SKP1-CUL1-F-box), an E3 ligase functioning in ubiquitination (Lv et al., 2017). Targets of MAX2 include SUPPRESSOR OF MORE AXILLARY GROWTH2-LIKE6 (SMXL6), SMXL7 and SMXL8 (Wang et al., 2015).

Stomata play a central role in carbon assimilation and stress responses as they regulate the uptake of CO_2_ which is inevitably connected to the evaporative loss of water. Moreover, open stomata provide an entry point for air pollutants and plant pathogens (Vainonen and Kangasjärvi 2015, Melotto et al., 2017). Guard cells which form the stomatal pore, respond to various endogenous and environmental stimuli by regulating their volume that in turn has a direct impact on the aperture of stomatal pores. Stomatal closure is induced by abscisic acid (ABA), pathogen-associated molecular patterns (PAMPs), high carbon dioxide (CO_2_) concentration, darkness, drop in relative air humidity and air pollutants such as ozone (Melotto et al., 2006, Merilo et al., 2013). ABA biosynthesis starts from carotenoids and ABA2 (ABSCISIC ACID DEFICIENT2) catalyses the conversion of xanthoxin to abscisic aldehyde (González-Guzmán 2002). Analysis of the *aba2* mutant that express *ABA2* with either guard cell- or phloem-specific promoter show that both promoters could restore ABA levels and functional ABA responses, demonstrating effective transport of ABA between tissues (Merilo et al., 2018). ABA-induced stomatal closure is initiated after binding of the hormone by PYR/PYL/RCAR receptors leading to inactivation of PP2C phosphates and followed by release of SNF-related protein kinases (SnRK2s) such as OST1 (OPEN STOMATA1). OST1 together with Ca^2+^-dependent protein kinases activate SLAC1 (SLOW ANION CHANNEL1) leading to stomatal closure (Merilo et al., 2013). Another protein, GUARD CELL HYDROGEN PEROXIDE-RESISTANT1 (GHR1), is required for stomatal closure and is proposed to act as a scaffold bringing together various proteins needed to initiate stomatal closure (Hua et al., 2012, Sierla et al., 2018).

We previously showed that the strigolactone perception mutant *max2* has increased susceptibility to plant pathogenic bacteria (*Pseudomonas syringae*) as a result from more open stomata and impaired stomatal closure in response to infection (Piisilä et al., 2015). Further, we demonstrated that the *max2* mutant also exhibits other stress-related phenotypes such as decreased tolerance to apoplastic reactive oxygen species (ROS), changes in stress-related gene expression, and hormonal signalling i.e. increased salicylic acid levels (Piisilä et al., 2015). However, as MAX2 is known to participate in several signalling pathways and acts as a central regulator in both strigolactone and karrikin signalling (Li et al., 2017), we set out to clarify the role of strigolactones in plant defence responses by analysis of strigolactone biosynthesis mutants (*max3, max4*) and their receptor (D14). To this end, we analysed the role of the strigolactone pathway in pathogen sensitivity, defence to ROS and stomatal regulation using single and double mutants defective in various steps of strigolactone biosynthesis and perception. Furthermore, the possible interaction between ABA and strigolactone signalling was assessed with a new set of double mutants.

## Materials and methods

### Plant material

All mutants used in this study were in the Col-0 genetic background. The following mutants were obtained from the Nottingham Arabidopsis Stock Centre: *max2-1* (Stirnberg et al., 2002), *max2-4* (SALK_028336), *max3-9* (Booker et al., 2004), *max3-11* (SALK_023975), *max4-1* (Sorefan et al., 2003) and *max4-7* (SALK_082552), *ost1-3* (*srk2e*, SALK_008068; Yoshida et al., 2002), *aba2-11* (González-Guzmán et al., 2002), *d14-1* (CS913109; Waters et al., 2012), *sid2-2* (Wildermuth et al., 2001). The *d14-seto5* and *max2-1 d14-seto5* (Chevalier et al., 2014) were obtained from Pilar Cubas. The *ghr1-3* (GK_760C07; Sierla et al., 2018) was donated by Jaakko Kangasjärvi. All mutants were genotyped by PCR based markers (Supporting Information Table 1).

The double mutants *aba2-11 max2-4, ghr1-3 max2-4* and *ost1-3 max2-4* were generated by crossing the respective single mutants with *max2-4* (pollen donor). Double homozygous plants were identified in F2 segregating progenies by PCR with gene-specific primers (Supporting Information Table 1).

### Growth conditions

Growth conditions in University of Tartu (the gas-exchange experiments). Arabidopsis seeds were planted in a soil pot covered by a glass plate with a hole through which the plants were grown as described by Kollist et al., (2007). The soil mixture contained 2:1 peat:vermiculite. Plants were grown in growth chambers in a 12 h photoperiod, 23°C/18°C temperature, 150 µmol m^−2^ s^−1^ light, and 70% relative humidity.

Growth conditions in University of Helsinki (all other experiments). Seeds were sown on a 1:1 peat:vermiculite mixture, vernalized in the dark for two days at 4°C, and germinated for one week. Next, plants were transferred to fresh pots to grow individually. Plants were grown in a growth room in 12 h light period (220 µmol m^−2^ s^−1^), 23°C/18°C day/night temperature and 70 % relative humidity. One to two weeks before the experiment plants were moved into a growth chamber with similar temperature/light conditions.

For gene expression experiments with plants treated with GR24, seeds were surface sterilized with 70 % ethanol and 2 % Triton X-100, rinsed 3 times with 99 % ethanol and dried on a filter paper. Plants were grown *in vitro* on ½ MS plates for 10 days in 16 h light/8 h dark cycle (110 µmol m^−2^ s^−1^), 23°C/18°C day/night temperature.

### Pathogen assays

*Pseudomonas syringae* pv. *tomato* DC3000 were grown in King’s B media at 28 °C overnight and the bacterial cells were collected by centrifugation at 6000 rpm for 8 min, and washed with 10 mM MgCl_2_. The centrifugation was repeated, and the bacteria were suspended in 10 mM MgCl_2_. OD_600_ was adjusted to 0.1, which equals to 10^7^ cfu ml^−1^ of bacteria in spray medium. To reduce surface tension 0.02 % Silwet L-77 was added just before inoculation. Next, 4-5 weeks-old plants were sprayed equally until their leaves were saturated with the spray medium. After inoculation plants were covered with plastic to maintain the humidity. The amount of bacterial cells was determined at 1.5 h and 48 h post inoculation. For each biological replicate, three leaf discs from three separate leaves were analysed. Leaf discs were surface sterilized with 70 % ethanol, washed with MQ water and ground in 0.2 ml of 10 mM MgCl_2_, after which the volume was adjusted to 1 ml. From the dilution series the aliquots of different dilutions were pipetted to King’s B media plates and grown for 2 days at 28 °C.

### Measurement of free SA levels

Free salicylic acid was measured using a modified biosensor-based method based on a protocol described by DeFraia et al., (2008). The bacterial biosensor strain *Acinetobacter* sp. ADPWH*_lux* (Huang et al., 2005, Huang et al., 2006) was grown overnight in LB medium at 37°C, after which the culture was diluted 1:20 and grown to an OD_600_ of 0.4. Leaf samples of 30 day-old plants were collected 27 h post spray inoculation with *Pst* DC3000 (OD_600_ = 0.2). A leaf disc (9 mm diameter) was cut from the 5th, 6th and 7th leaf of each plant and the three discs were homogenized in 200 µl of LB-medium. To count bacterial cfu, we took 8 µl aliquot, and the rest of the sample was centrifuged for 15 min at 21000g. Next, 20 µl of the supernatant was mixed with 30 µl of LB-medium and 50 µl of 1:100 diluted biosensor culture (OD_600_ = 0.004) on a white 96-well plate. The plate was incubated at 37 °C for one hour without shaking, after which the luminescence was measured with Perkin Elmer EnSpire 2300 plate reader. For standard curve, 0, 5, 10, 15 and 20 ng of sodium salicylate in 30 µl of LB-medium were mixed with 20 µl of plant extract from *sid2-2* plants and 50 µl of the diluted biosensor strain. The standard curve samples were measured at the same time as the samples. Due to non-linearity of the standard curve values, separate linear best-fit models were fitted for low (0-10 ng) and high (10-20 ng) amounts of salicylic acid standard as described by (DeFraia et al., 2008). The luminescence values of the samples were converted to estimated masses of salicylic acid based on the standard curves and reported as ng cm^−2^. Statistical analysis was performed in R programming environment and figures were prepared with ggplot2 package in R (R Core Team 2018, Wickham 2016).

### Ozone exposure and ion leakage analysis

The ozone exposure was conducted with 350 nL L^−1^ of ozone gas for 6 hours on 4-5 week old plants as described by Overmyer et al., 2000. The relative ion leakage was measured as conductivity in 0, 6 and 24 hours after beginning of ozone exposure. There were four biological repeats for each time point from each plant line. The conductivity was measured using Horiba Twin Cond Conductivity Meter B-173.

### Expression analysis by qPCR

*Pseudomonas syringae* pv. *tomato* DC3000 bacteria were grown as described previously and infection was done with OD_600_ = 0.1 on 4-5 week old plants. The samples were taken 0 (i.e. non-infected), 3, 6, 24 and 48 h after the infection.

GR24 (Chiralix) was dissolved in DMSO and a solution of 10 µM GR24 and 0.01 % of Silwet L-77 was poured into the wells of a 6-well multi plate. The control solution consisted of the corresponding amounts of DMSO and Silwet L-77. Whole intact 10-day-old in vitro grown Col-0 seedlings were put into the wells and samples were collected after 3 hours.

RNA was isolated with GeneJET Plant RNA Purification Mini Kit (Thermo Scientific). RNA (3 µg in GR24 assay/2 μg in Pseudomonas infection) was DNAseI treated and reverse transcribed with Maxima RT and Ribolock Rnase inhibitor (Thermo Scientific) according to the manufacturer instructions. After cDNA synthesis the reactions were diluted to a final volume of 100 µl. qPCR was performed in triplicate using 5× HOT FIREPol EvaGreen qPCR Mix Plus (Solis Biodyne). The cycle conditions with Bio-Rad CFX384 were: 1 cycle initiating with 95°C 10 min, 50 cycles with 95°C 15s, 60°C 30s, 72°C 30s and ending with melting curve analysis. Normalization of the data was performed in qBase 3.0 (Biogazelle), with the reference genes PP2AA3, TIP41 and YLS8 in GR24 assay / F-box protein AT5G15710 in Pseudomonas infection. Primer amplification efficiencies were determined in qBase from a cDNA dilution series. All primers are listed in Supporting Information Table 1.

### Stomatal conductance

The porometer measurements were done on 5-6 week old plants with an AP4 porometer (Delta-T Devices, Cambridge, UK). The plants were measured according to the phenotype-based growth stage (Boyes et al., 2001) and 2-3 leaves per each growth stage was measured from each plant and altogether minimum 20 plants were measured per each plant line. The age of the plants was 5-6 weeks when the rosettes reached their maximal size but the flower buds were not yet visible.

The basal level of whole-plant stomatal conductance and stomatal responses to CO_2_, darkness and 5 µM ABA foliar spray were measured with a custom-made gas exchange measurement device at the University of Tartu as described in details by Kollist et al., 2007. The unit of stomatal conductance mmol m^−2^ s^−1^ reflects the amount of H_2_O moles that exits the plant through stomata per m^2^ of leaf area per second. The age of the plants was 3-4 weeks when they were measured. Due to the bushy phenotype of several studied mutants, the rosette area of all plants was calculated by separating the leaves and measuring individually. Prior to the experiment, plants were acclimated in the measurement cuvettes at ambient CO_2_ concentration (∼400 ppm), 100 μmol m^−2^ s^−1^ light, and ambient humidity (RH 65%–80%) for at least 1 h or until stomatal conductance was stable.

ABA-induced stomatal closure was induced by foliar spray with 5 μM ABA solution (5 μM ABA, 0.012% Silwet L-77, 0.05% ethanol). At time point T=0 min, plants were removed from the measuring cuvettes and sprayed with either 5 μM ABA solution or control solution (0.012% Silwet L-77, 0.05% ethanol). Thereafter, plants were returned to the cuvettes and stomatal conductance was monitored for 40 min.

Foliar spray with GR24 was conducted as following: at time point T = 0 min acclimated plants were removed from the measuring cuvette and 10 μM GR24 solution (10 µM GR24, 0.012% Silwet L-77 solution, 0.02% DMSO) was applied with a spray bottle 5 times approx. 30 cm from the rosette so that the plant would look slightly wet. Next, plants were returned to the measuring cuvettes to dry and starting from T = 8 min stomatal conductance was monitored for 56 minutes.

Statistical analyses of gas exchange data were performed with Statistica, version 7.1 (StatSoft Inc., Tulsa, Oklahoma, United States). All effects were considered significant at p < 0.05.

### Accession numbers

ABA2 - AT1G52340, MAX2 – AT2G42620, OST1 - AT4G33950, GHR1 - AT4G20940, MAX3 - AT2G44990, MAX4 - AT4G32810, D14 - AT3G03990, ICS1/SID2 - AT1G74710, PR1 - AT2G14610, FRK1 - AT2G19190, NCED3 - AT3G14440, GRX480 - AT1G28480, AXR3/IAA17 - AT1G04250, PP2AA3 - AT1G13320, YLS8 - AT5G08290, TIP41- AT4G34270.

## Supporting Information

Supporting Information Figure 1. The sensitivity to pathogens in darkness.

Supporting Information Figure 2. Growth of *P. syringae* in free SA analysis experiment.

Supporting Information Figure 3. The time course data used to calculate the bar graphs of Figure 7(c)–7(h).

Supporting Information Figure 4. The rosette morphology of Arabidopsis Col-0 and the mutant plants used in this study.

Supporting Information Table 1. The PCR primers used in genotyping and qPCR primers used in gene expression analysis.

## Results

### Strigolactone affects sensitivity to pathogens in Arabidopsis

Plant defence against pathogens involves a network of interacting signalling pathways where several plant hormones are key components (Overmyer et al., 2018). Recently, strigolactones were identified as an additional component that regulates drought- and pathogen responses (Bu et al., 2014, Ha et al., 2014, Piisilä et al., 2015). Here, we further explored the mechanism of strigolactones in sensitivity to pathogens by analysis of strigolactone receptor (D14) mutants and strigolactone synthesis (MAX3 and MAX4) mutants in order to find biological processes regulated by strigolactones.

Spray infection with *P. syringae* DC3000 allows quantification of pathogen sensitivity in relation to stomatal immunity, as the bacteria need to enter the plant through stomata to multiply (Melotto et al., 2006). Therefore, we investigated the pathogen sensitivity of strigolactone synthesis and perception mutants. Interestingly, 1.5 hours post inoculation (hpi), only *max2* mutants exhibited increased sensitivity to *P. syringae* DC3000 infection (Figure 1; OD_600_ = 0.1) compared to Col-0. However, at a later time point (48 hpi) strigolactone sensing (*max2*), the receptor (*d14*) and synthesis (*max3* and max*4*) mutants were all more sensitive to *P. syringae* DC3000 (Figure 1). Moreover, at 48 hpi some differences were observed between the mutants, notably the *max2-4* allele (a SALK T-DNA line) had a stronger phenotype than an EMS mutant *max2-1* (Stirnberg et al., 2002). Similarly, the Wisconsin T-DNA knock-out line *d14-1* had a stronger phenotype than an EMS mutant *d14-seto5* (Chevalier et al., 2014). In order to assess the role of stomatal openness in the infection, a similar experiment was also done in an inverted light rhythm i.e. the plants were infected in darkness when the stomata are normally closed, but the result was rather similar to the one obtained when infection was performed in normal light conditions (Supporting Information Figure 1). Therefore, we conclude that both strigolactone biosynthesis and perception mutants were all significantly more sensitive to *P. syringae* DC3000 spray infection than Col-0 at 48 hpi.

**Figure 1.**
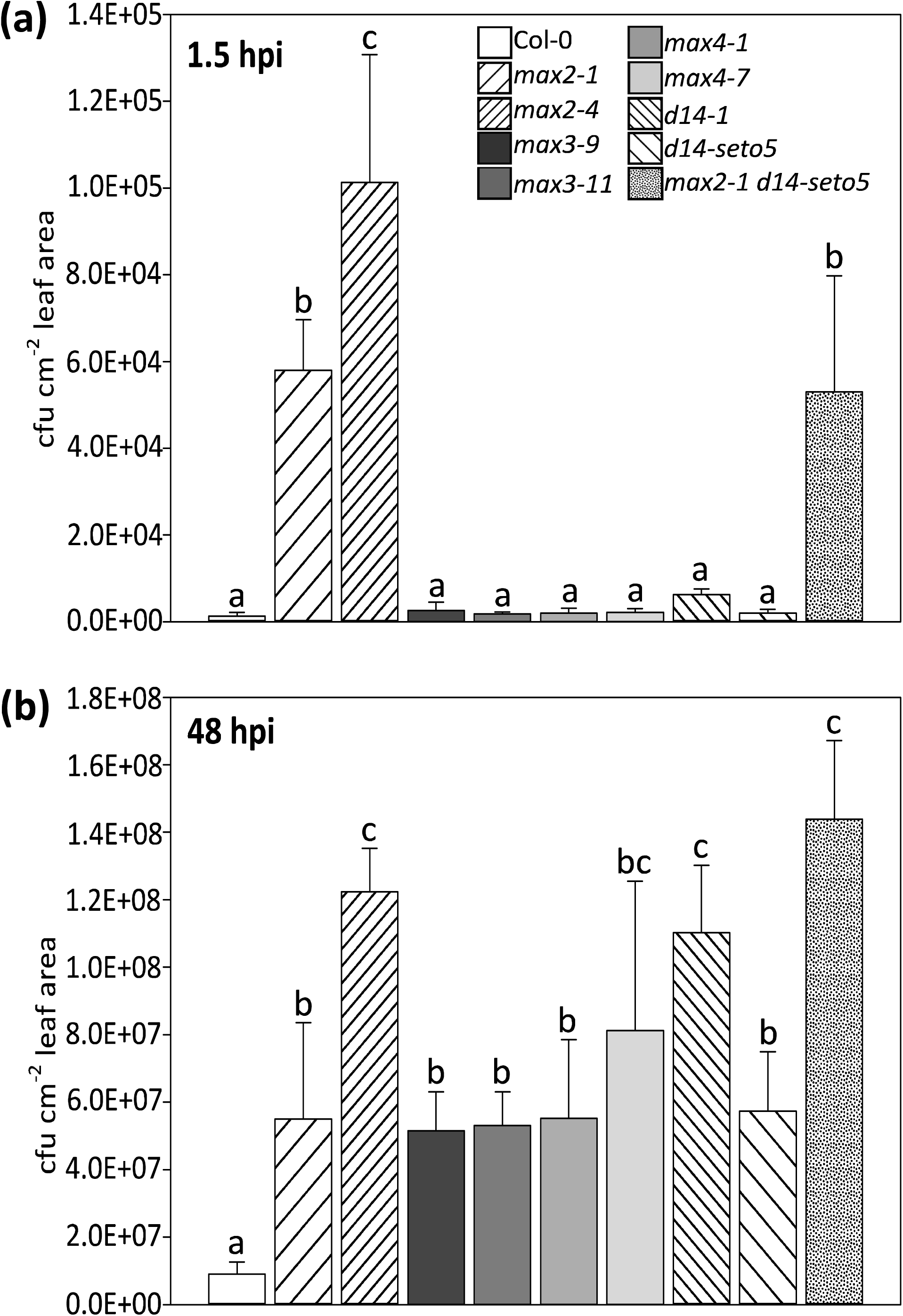
Bacterial calculation after infection with *Pseudomonas syringae* DC3000. The infection was done by spraying followed by calculation of bacteria at 1.5 hours and 48 hours post inoculation. In each experiment four plants per line and three leaves per plant were used to measure the bacterial concentration. The experiment was repeated three times with similar results. The results are shown as means ± SE. In statistical analysis we compared if Col-0 significantly differs from mutants; first a logarithmic transformation was conducted on the data, then univariate analysis of variance combined to Hochberg post hoc test.

### *max2* mutants accumulate high level of free salicylic acid in response to *P. syringae* DC3000

The plant hormone salicylic acid (SA) has a key role in defence against pathogen infection (Fu and Dong 2013, Pieterse 2012). Therefore, we measured free SA levels in Col-0 and the strigolactone biosynthesis/perception mutants at 27 hpi after spray infection with *P. syringae* DC3000 (Figure 2; OD_600_ = 0.2, which was a higher inoculum than the one used in Figure 1; OD_600_ = 0.1). Simultaneously, we measured the extent of pathogen growth at the same time point (Supporting Information Figure 2). As a control for the assay, we included the SA biosynthesis mutant *sid2*, which does not accumulate SA in response to pathogens (Wildermuth et al., 2001). High levels of SA accumulated in both *max2* mutants (Figure 2), consistent with our previous measurements of SA after pathogen infection (Piisilä et al., 2015). In contrast, the levels of SA did not increase to significantly higher levels in either strigolactone biosynthesis mutants (*max3, max4*) or the receptor (*d14*). At this higher pathogen inoculum and earlier time point (27 hpi vs 48 hpi), only the *max2* mutants were significantly more sensitive (Supporting Information Figure 2). Therefore, we conclude that the *max2* mutants are pathogen sensitive at an earlier time point also when a higher inoculum of bacteria is used, but the differences in sensitivity of strigolactone biosynthesis and perception mutants were only significant with smaller bacterial inoculum at a later time point.

**Figure 2.**
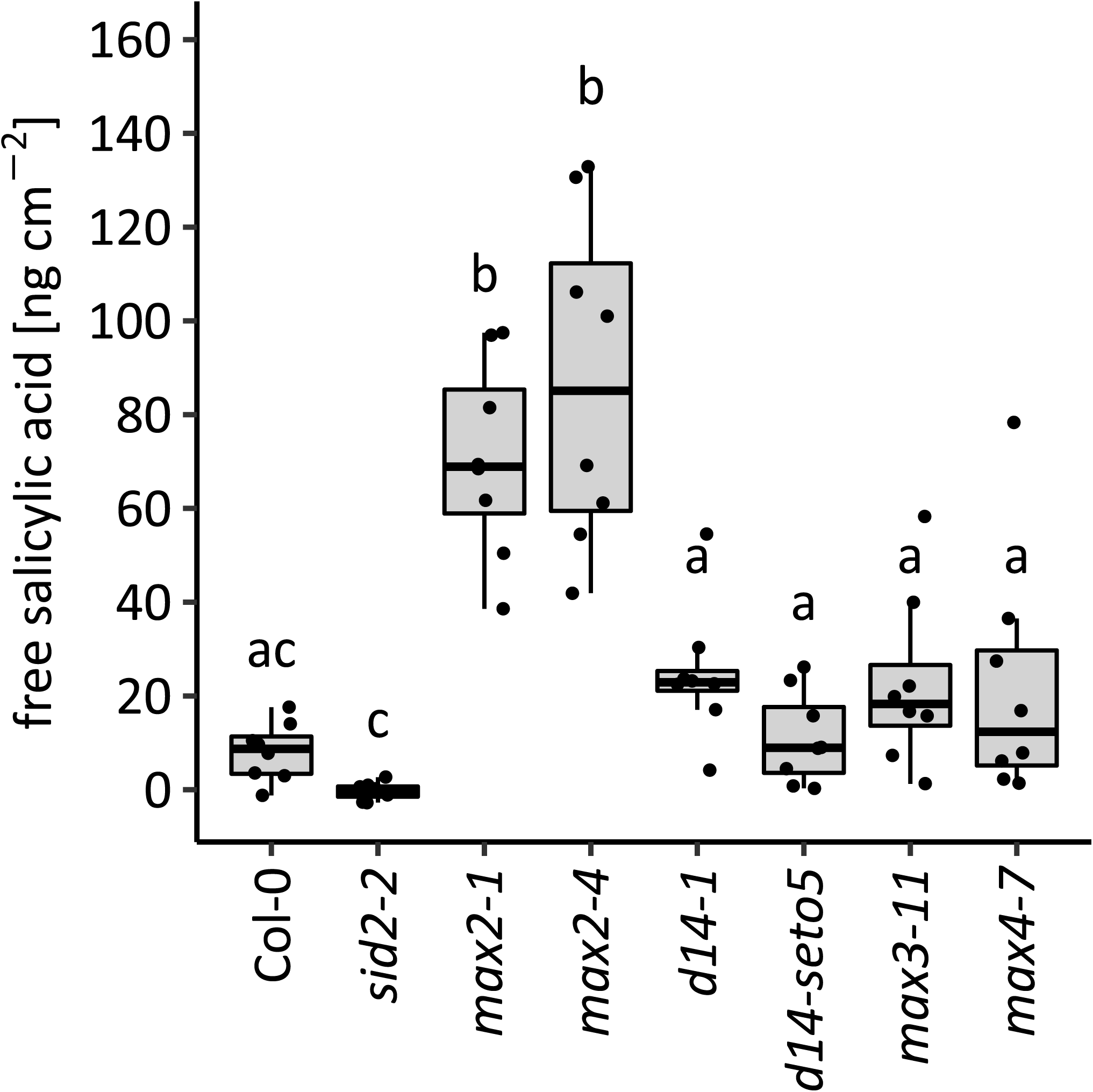
Accumulation of free salicylic acid in response to *Pseudomonas syringae* DC3000. Wild type (Col-0), strigolactone signalling mutants (*max2-1, max2-4, d14-1* and *d14-seto5*) and strigolactone biosynthesis mutants (*max3* and *max4*) were infected with *P. syringae* (OD_600_ = 0.2) by spraying. SA was measured using a biosensor-based method based on the protocol by DeFraia et al., (2008). Salicylic acid biosynthesis mutant (*sid2-2*) was included as a control for accumulation of free salicylic acid. Salicylic acid accumulation was measured 27 hours post inoculation. Each dot represents one plant. In total, seven to eight plants were used per line. The experiment were performed three times with similar results. Box plots are summarising data by showing the median, and first and third quartiles. Whiskers are extending to a maximum of 1.5 × interquartile range beyond the box. Different letters indicate significant differences (P < 0.05) as determined by Kruskal-Wallis Rank Sum test followed by Pairwise Wilcoxon Rank Sum tests with multiple testing correction to p-values using Holm method.

### Strigolactone is not a major regulator of defence gene expression

The role of strigolactones in Arabidopsis is best described in branching (Gomez-Roldan et al., 2008, Goulet and Klee 2010, Crawford et al., 2010, Waldie et al., 2014). Microarray analysis to find strigolactone-regulated genes was previously performed with GR24 (a strigolactone analogue) (Mashiguchi et al., 2009). In that study, 64 genes had significantly altered expression and the magnitude of transcriptomic response (i.e. fold change) in the GR24-responsive genes was small (Mashiguchi et al., 2009). To test if GR24 can regulate genes related to pathogen defence, we treated 10-days-old in vitro grown Col-0 with 10 µM GR24 for 3 h and measured gene expression with real-time quantitative PCR (qPCR). Expression of the well-established pathogen-responsive genes *PR1* (*PATHOGENESIS-RELATED GENE 1*, a late response gene indicating activated SA response, Uknes et al., 1992) and *FRK1* (*FLG22-INDUCED RECEPTOR-LIKE KINASE 1*, an early flg22 response marker gene, Asai 2002) decreased, although the differences were not statistically significant. Consistent with the role of strigolactones acting together with auxin in regulation of plant development, expression of *AXR3/IAA17 (AUXIN RESISTANT 3/ INDOLE-3-ACETIC ACID INDUCIBLE 17)* increased (Figure 3). Expression of *GRX480* (a glutaredoxin that regulates protein redox state) also increased. Notably, *GRX480* expression is regulated by several stimuli, including SA and ROS (Blanco et al., 2009, Koornneef and Pieterse 2008, Pieterse et al., 2012, Xu et al., 2015a).

**Figure 3.**
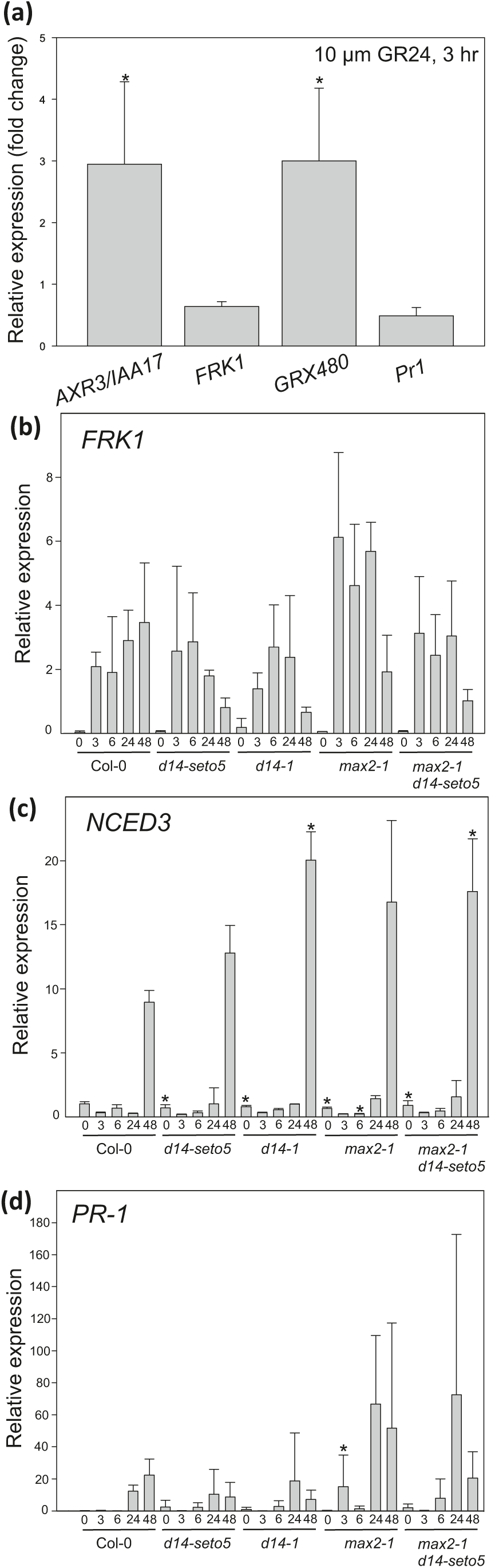
Relative gene expression in response to *Pseudomonas syringae* DC3000 spray infection and GR24 spray (dissolved in DMSO). (a) Relative expression (fold change GR24/control) after 3 h 10 µM GR24 treatment. The fold change is calculated from three biological repeats. (b - d) Four week old plants were spray infected with *P. syringae* DC3000. Relative expression was calculated from three biological replicates for each plant line in each time point. In statistical analysis we compared if Col-0 significantly differs from mutants. We conducted a logarithmic transformation on the data, then univariate analysis of variance combined to the Hochberg post hoc test.

We also monitored the expression of *PR1, FRK1* and *NCED3* (*NINE-CIS-EPOXYCAROTENOID DIOXYGENASE 3*, a gene encoding a protein that is catalysing a major step of the ABA biosynthesis pathway (Fan et al., 2009, Tan et al., 2003, Wang et al., 2011)) in *P. syringae* infected *max2* and *d14* plants. A few subtle differences were observed including increased expression of PR-1 at an early time point (3 hpi) in *max2-1* and increased expression of *NCED3* at the start (0 hpi) and end (48 hpi) of the experiment (Figure 3). As a relatively large variation was observed between different biological repeats, it is possible that some subtle differences between mutants might be obscured, but overall it appears that strigolactone signalling is not a major regulator of defence-related genes.

### Strigolactone perception mutants, but not biosynthesis mutants are ozone sensitive

Treatment of plants with ozone serves as a method to explore plant sensitivity to apoplastic ROS. Ozone enters plants via stomata and immediately degrades to ROS (O_2_ ^•-^ and H_2_O_2_) in the apoplastic space, which initiates cell death signalling and leads to development of visible tissue lesions (Overmyer et al., 2018, Vahisalu et al., 2010, Vainonen and Kangasjärvi 2015). Importantly, as ozone enters the plant through stomata, the mechanisms of sensitivity to this air pollutant can broadly be divided into stomata-dependent or -independent mechanisms (Xu et al., 2015b, Vainonen and Kangasjärvi, 2015). Therefore, we assessed the extent of ozone-induced damage observed after a 6 h treatment with 350 nL L^−1^ O_3_ by measuring ion leakage before ozone exposure (0h) and at two time points (6 h and 24 h) after beginning of the treatment (Figure 4). Similarly to phenotypes observed after infection with *P. syringae* (Figure 1a and Supporting Information Figure 2), only the *max2* mutants were ozone sensitive. As also seen in the pathogen infection, the *max2-4* allele had a stronger phenotype than *max2-1*. The ion leakage measured 24 hours after beginning of exposure was also higher in *max2-1 d14-seto5* double mutant as compared to *max2-1* or *d14-seto5* single mutants. The differential response of *max2* versus the biosynthesis and perception mutant suggest that MAX2 might act in additional signalling pathways that extend beyond strigolactone signalling.

**Figure 4.**
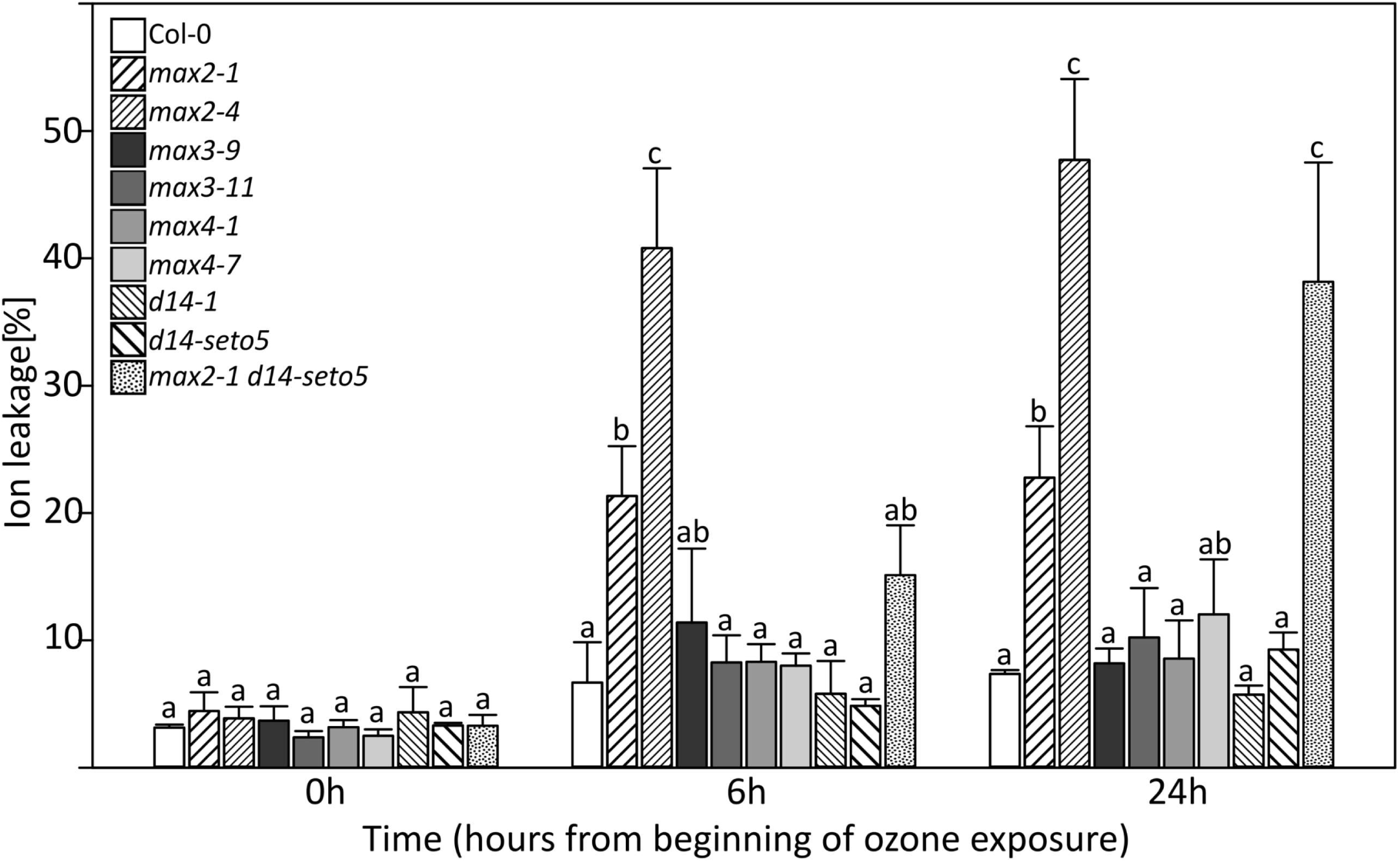
Ozone sensitivity of strigolactone biosynthesis and perception mutants. Ion leakage was measured at indicated time points counted from the beginning of ozone exposure (350 nL L^−1^ for 6 hours). In statistical analysis we conducted a logarithmic transformation on the data, then univariate analysis of variance combined to Hochberg post hoc test. The ion leakage was calculated as percentage of conductance related to the amount of total ions in the sample. The experiment was repeated three times and there were 4 biological repeats for each plant line in each time point.

### Strigolactones do not directly regulate the stomatal aperture

Strigolactone perception and biosynthesis mutants were more sensitive to *Pseudomonas* infection (Figure 1). However, changes in SA accumulation (Figure 2) or gene expression (Figure 3) could not explain this sensitivity. As spray infections require the bacteria to enter through stomata, guard cell signalling leading to stomatal closure may be one of the primary functions of strigolactones. To explore this possibility we used various stomatal assays and mutants to further study strigolactone-dependent regulation of guard cell function (Figures 5 – 8). First, we tested the ability of GR24 to induce rapid stomatal responses within an hour after application onto leaves. For this we sprayed the Col-0 rosettes with 10 µM GR24 (stock dissolved in DMSO and diluted in water) and followed the stomatal conductance by measuring the whole-rosette stomatal conductance (Kollist et al., 2007, Merilo et al., 2015). However, no change in stomatal conductance after the foliar spray was detected neither in mock-nor GR24-treated plants (Figure 5a). Next, we tested the ability of GR24 to close stomata in a longer time frame. GR24 is not soluble in water, therefore, alternative solvents have been used to dissolve this chemical (DMSO was used for gene expression, Mashiguchi et al., 2009; acetone was used for stomatal aperture measurements, Lv et al., 2017). We prepared a stock of GR24 in DMSO or acetone and diluted to 5 µM in water with 0.02 % Silwet L-77 and sprayed on Col-0 plants. Stomatal apertures were measured 24 hours after spraying with the use of a protocol formulated by Chitrakar and Melotto (2010). This late time point was selected to identify possible long-term effects of GR24 on stomatal aperture. However, no significant effect of GR24 on stomatal aperture was observed (Figure 5b). Taken together, our results indicate that treatments with a strigolactone analogue GR24 are not able to induce stomatal closure in intact plants.

**Figure 5.**
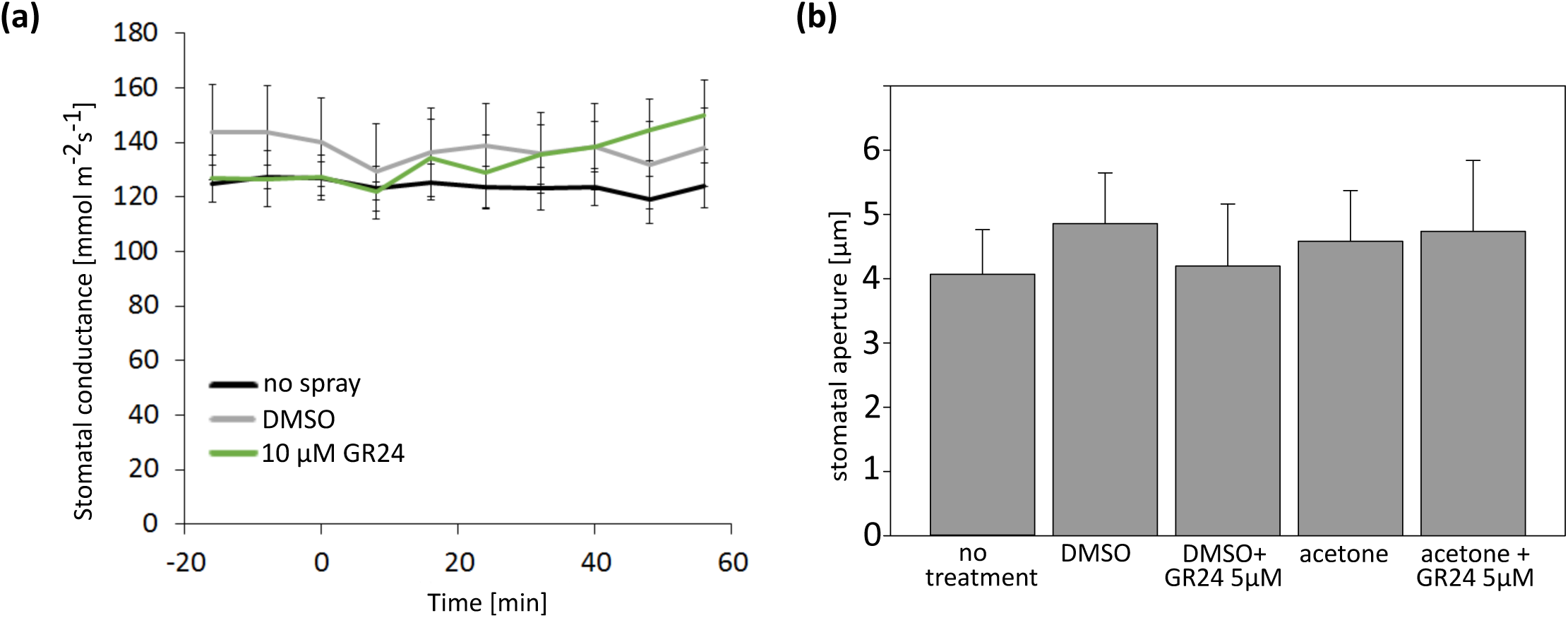
Stomatal response to strigolactone analogue (GR24) spray. (a) Time course of stomatal conductance of WT Col-0 plants. At time T=0 min the plants were sprayed with 10 µM GR24 or mock and returned back to the measuring cuvette. Data is presented as a mean ± SEM (n=5). (b) The stomatal aperture width in response to GR24 stock dissolved in DMSO or acetone. The Col-0 plants were sprayed fully wet with 5 µM GR24 or an equivalent mock solution a day before the stomatal aperture measurements and the differently treated plants were kept covered separately with plastic overnight. Data is presented as a mean ± SE. Altogether approximately 200 stomata were measured from leaves of five sprayed plants, and the experiment was repeated three times with similar results.

### Stomatal conductance is higher in strigolactone mutants at different leaf developmental stages

Various types of methods are available to measure stomatal conductance and thus getting an estimate of stomatal aperture. To investigate the stomatal conductance at the developmental stage-resolution we measured stomatal conductance with a porometer, in which the sensor head is clipped onto a single leaf and the conductance is measured from only the abaxial side of the leaf. For this, we used 5-6 weeks-old plants and measured leaves at different developmental stages 1-2, 3-4 and 5 (Boyes et al., 2001, which roughly corresponds to young, middle-age and old, leaves respectively, Figure 6). Most of the strigolactone perception and biosynthesis mutants had higher stomatal conductance as compared to Col-0 with no regard to leaf age (Figure 6). The weak *d14-seto5* did not display higher conductance, and the double mutant *max2-1 d14-seto5* showed a differential response with increased conductance only in middle age and older leaves (Figure 6).

**Figure 6.**
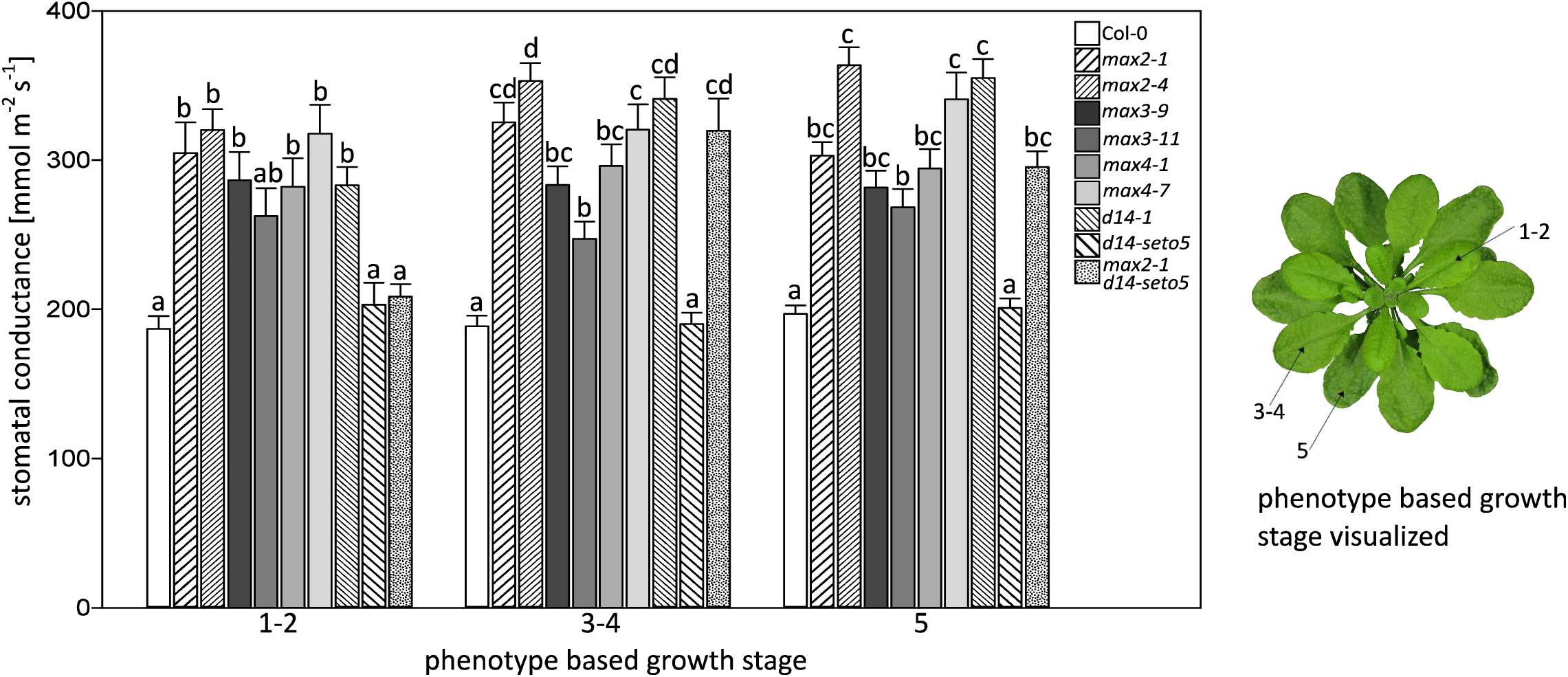
The stomatal conductance of strigolactone biosynthesis and perception mutants measured with a porometer from leaves of different developmental stages. The error bars represent standard error of the mean. 2-3 leaves per each growth stage was measured from each plant and altogether a minimum 20 plants were measured from each plant line. The phenotype based growth stage is determined in the article by Boyes et al., (2001) in which numbers indicate the growth stage; 1 indicates leaf production, 3 rosette growth and 5 inflorescence emergence. We used the plants for analysis before they reached the stage 5.10 (i.e. before the first flower buds were visible). In statistical analysis we conducted a logarithmic transformation on the data, then univariate analysis of variance combined to Tukey HSD post hoc test.

### Stomatal conductance in intact plants and in response to darkness, high CO_2_ and ABA

Porometer measurements requires clipping the sensor head onto the leaf, which could potentially activate stress responses e.g. touch-induced signalling. To measure the stomatal conductance of intact whole plant rosettes, we used a multi-cuvette gas exchange system (Kollist et al., 2007), in which the plant is inserted into the machine without any touching. In contrast to results obtained with the porometer (Figure 6), the whole-rosette stomatal conductivity measurements indicated that only the *max2* alleles (and *max2-1 d14-seto5*) displayed increased stomatal conductance (Figures 7a and 7b).

**Figure 7.**
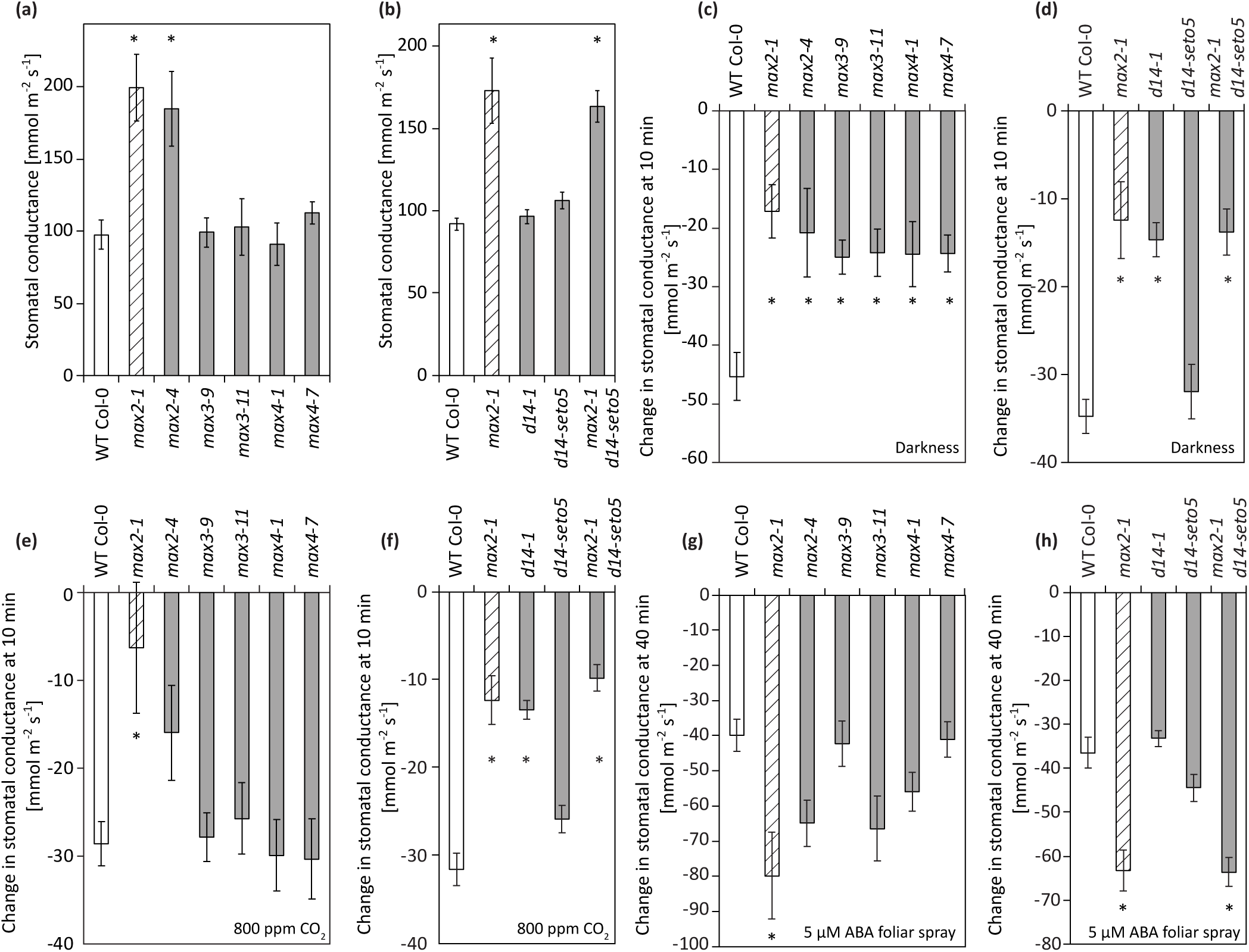
Whole plant stomatal conductance and stomatal closure responses to darkness, high CO_2_ and ABA foliar spray. (a) Stomatal conductance of *max2, max3* and *max4* mutants (n= 7-11). (b) Stomatal conductance of *max2* and *d14* mutants (n = 10-20). (c) Darkness-induced stomatal closure of *max2, max3* and *max4* mutants (10 min after induction; n = 7-11). (d) Darkness-induced stomatal closure of *max2* and *d14* mutants (10 min after induction; n = 10-20). (e) High CO_2_-induced stomatal closure of *max2, max3* and *max4* mutants (10 min after induction; n = 6-12). (f) High CO_2_-induced stomatal closure of *max2* and *d14* mutants (10 min after induction; n = 10-23). (g) ABA-induced stomatal closure of *max2, max3* and *max4* mutants (40 min after induction; n = 6-12). (h) ABA-induced stomatal closure of *max2* and *d14* mutants (40 min after induction; n = 13-23). All graphs present the mean ± SEM. Asterisks denote statistically significant differences according to one-way ANOVA with Tukey HSD post hoc test. The time course data used to calculate the bar graphs (c)–(h) can be found in Supporting Information Figure 3.

Next we measured the response to several other stimuli: darkness, high CO_2_ and ABA (Figure 7) that induce stomatal closure in wild-type plants (Assmann and Jegla 2016, Merilo et al., 2013). In response to darkness, both strigolactone biosynthesis (*max3, max4*) and perception (*max2, d14-1*) mutants had significantly slower rate of stomatal closure while upon high CO_2_ treatment the response was impaired in *max2* and *d14* mutants (Figures 7e and 7f). In contrast, the stomatal response to ABA was not impaired in any of the mutants, instead an enhanced stomatal closure rate was observed in *max2-1*, possibly due to more open stomata of *max2-1* before ABA treatment (Figures 7g and 7h).

Most of our knowledge on regulation of guard cell signalling is in the context of ABA signalling (Merilo et al., 2018), and very few regulators specific for other stimuli have been found (Engineer et al., 2016). Thus the differential response of especially *max2* with a defective response towards high CO_2_ and darkness, and normal to enhanced response towards ABA, adds a new regulatory layer in guard cell signalling. As an F-Box protein, MAX2 might be involved in the targeted degradation of some other regulatory component in guard cell signalling.

### MAX2 regulates stomatal function independently of ABA signalling

Multiple results obtained in this study indicated significant differences between *max2* and other strigolactone signalling-related mutants i.e. the *max2* mutant was the only mutant exhibiting ozone sensitivity and consistently higher stomatal conductance (Figures 4, 6 and 7). To further explore the function of MAX2 in the guard cell signalling network, we crossed *max2* with mutants defective in ABA biosynthesis (*aba2*), guard cell ABA signalling (*ost1*) and a scaffold protein GHR1 (*ghr1*) that is required for activation of the guard cell anion channel SLAC1. Next, single and double mutants were subjected to measurement of stomatal conductance. Interestingly, all double mutants had significantly higher stomatal conductance than the corresponding single mutants (Figure 8), suggesting that MAX2 acts in a signal pathway that functions in parallel to the well-characterized stomatal ABA signalling pathway.

**Figure 8.**
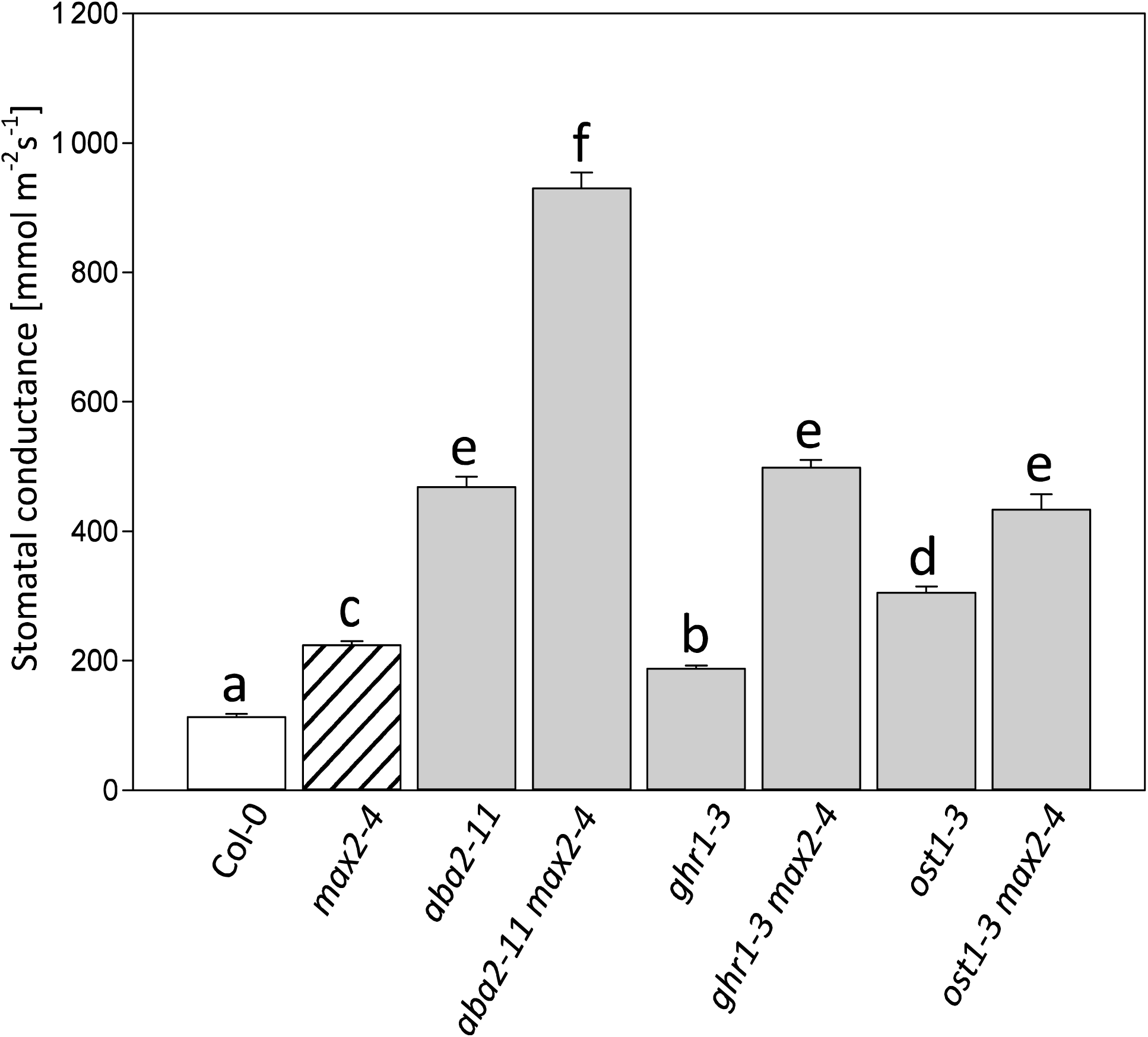
The basal level of whole-rosette stomatal conductance of double mutants measured with intact plants. Four weeks old plants were measured, and the stomatal conductance is the average from 5-6 plants. Data is presented as the mean ± SEM. In statistical analysis we conducted a logarithmic transformation on the data, then univariate analysis of variance combined to Tukey HSD post hoc test.

## Discussion

Plant defence responses to pathogen infection are highly complex and include many different signalling pathways (Koornneef and Pieterse 2008, Pieterse et al., 2012). While hormones associated with stress e.g. salicylic acid, jasmonic acid, ethylene and abscisic acid have long been studied for their role in pathogen responses, also hormones typically associated with development influence the outcome of plant-pathogen interactions (Pieterse et al., 2009). While the function of strigolactones in Arabidopsis was initially characterized for their role in shoot branching (Bennett et al., 2006, Hayward 2009, Stirnberg et al., 2007), they appear to be important in pathogen sensitivity. Mutants involved in strigolactone sensing (*max2, d14*) and biosynthesis (*max3* and *4*) were more sensitive to *P. syringae DC3000* spray infection than Col-0 (Figure 1b). However, given the differential response when higher inoculum was applied, where only *max2* showed increased sensitivity (Supporting Information Figure 2), it seems like the role of strigolactone is better described as a modulator of the defence response rather than a master regulator. Similarly the very modest response of pathogen-defence genes to the application of the synthetic strigolactone analogue GR24 (Figure 3 and Mashiguchi et al., 2009), also indicates that strigolactones are not direct regulators of defence signalling.

We propose that the role of strigolactone signalling components, especially MAX2, in plant defence, is related to the regulation of stomatal function, which subsequently influences pathogen and defense responses. This was clearly indicated by an increased ozone sensitivity of *max2* mutant plants (Figure 4). Arabidopsis mutants and accessions with more open stomata are ozone sensitive as more ozone can enter the plant (Overmyer et al., 2008, Brosché et al., 2010). Similarly more open stomata would also allow higher entry of pathogenic bacteria e.g. *P. syringae*. Additionally, the impaired stomatal closure in response to *P. syringae* infection likely contributes to sensitivity (Piisilä et al., 2015). In pathogen- and ozone-sensitivity studies, the strong *max2-4* allele was clearly more impaired than the weak *max2-1*. When weak alleles of *max2* (*max2-1*) and *d14* (*d14-seto5*) were combined in the *max2-1 d14-seto5* double mutant a similar phenotype to the strong *max2-4* was observed in pathogen (48 hpi) and ozone (24 h) responses (Figures 1 and 3). This is consistent with the current understanding of D14 and MAX2 interacting in strigolactone perception (Lv et al., 2017). However, the fact that *d14-1* and strigolactone biosynthesis mutants *max3* and *max4* were not ozone sensitive suggests that MAX2 has a more versatile role in stress responses than the other strigolactone signalling-related proteins. Previously MAX2 was also shown to have a role in karrikin signalling (Li et al., 2017), pointing towards different signalling roles for this F-box protein, possibly by targeting the degradation of proteins in different signalling pathways.

To further evaluate the role of strigolactones in regulation of guard cell signalling we used several different methods and genetic tools. To our surprise we could not observe GR24-induced stomatal closure in either stomatal aperture or stomatal conductance assays (Figure 5). Ha et al., (2014) rescued the drought phenotype of the strigolactone biosynthesis (*max3* and *max4*) mutants with strigolactone spray. Moreover, by using epidermal peels, Lv et al., (2017) showed that the stomata close in response to GR24, and strigolactones were concluded to be common regulators of stomatal closure *in planta* (Zhang et al., 2018). One challenge in using GR24 is that no standard method for dissolving this chemical is established and possibly the solvent of GR24 might affect the results. Thus, we dissolved GR24 both in DMSO and acetone but neither of the solutions resulted in clear differences from mock in the size of stomatal aperture or in stomatal conductance.

There are several methods to measure stomatal function. First we performed a classical porometer measurement in order to explore if the growth stage (or age) of the leaves affected stomatal conductance. All of the strigolactone biosynthesis and perception mutants had a higher stomatal conductance except for *d14-seto5* (Figure 6) that consistently had weaker phenotypes than the other mutants, which is also evidenced by its growth phenotype (Supporting Information Figure 4). As the strigolactone mutants have a “bushy” phenotype (Supporting Information Figure 4), with leaves often laying on top of each other, the porometer measurements have the advantage of measuring a specific area independent from the rest of the plant. However, the data provided by the porometer are also limited, since the porometer sensor area is rather small and thus only a small area of a leaf is measured and we only measured the abaxial side of the leaf.

To complement the porometer data we used a method that measures the whole-rosette stomatal conductance using a customized gas-exchange device which allows for measurement of intact plants and real-time responses to stomata-affecting factors, such as CO_2_ concentration, darkness and the phytohormone ABA (Kollist et al., 2007). In contrast to porometer measurements, only *max2* and *max2 d14* mutants had more open stomata when the intact whole rosettes were measured (Figures 7a and 7b.). The difference in results obtained with these two methods might have several explanations: (1) in promoter measurements only a limited leaf area can be measured and the measurement involves touching the plants (2) In whole-rosette measurements the “bushy” phenotype with overlapping leaves of the strigolactone signalling mutants might create a microclimate that alters stomatal conductance. (3) As porometer measurements were performed at the University of Helsinki, and the whole-rosette assays at the University of Tartu, other factors affecting growth conditions could sensitize the biosynthesis mutants (*max3, max4*) to have more open stomata in the Helsinki growth conditions. Further research might resolve this issue, but given the consistent phenotype of *max2* across several different assays and growth conditions, the response of this mutant strongly suggests an important role for MAX2 in guard cell signalling.

Testing stomatal responses to several different treatments showed that high CO_2_-induced stomatal closure was impaired in *max2* and *d14-1* (Figures 7e and 7f). A sudden darkness treatment during the normal light period is partially initiated by the same mechanism as CO_2_ signalling. Removal of light stops photosynthesis and this leads to increase of CO_2_ concentration inside the leaf, similar to the situation when elevated CO_2_ is applied. The darkness-induced stomatal closure was reduced in *max2, max3, max4* and *d14-1*; i.e the response to darkness was more broadly impaired than the response to high CO_2_. Of the different stimuli and treatments that lead to stomatal closure, the response to darkness might be the least studied. Thus, the impaired darkness response in both strigolactone biosynthesis and perception mutants opens the possibility for further studies into this branch of guard cell signalling.

To further study the relationship between strigolactone and ABA, we examined if ABA signalling and *MAX2* share the same elements in guard cell signalling. For this, we crossed *max2* with other guard cell signalling mutants *ost1, ghr1* and the ABA biosynthesis mutant *aba2.* The resulting double mutants (*max2 ghr1, max2 ost1* and *max2 aba2*) had a higher stomatal conductance than any of these mutants individually. Thus, it appears that MAX2 functions on a pathway that is parallel to ABA signalling (see also Lv et al., 2017). As very few regulators of ABA-independent guard cell signalling have been found (Assman and Jegla 2016, Engineer et al., 2016), the impaired CO_2_ and darkness response in *max2* implicate that the F-Box protein MAX2 has a crucial role in targeting as yet unidentified important guard cell regulator to ubiquitin-mediated protein degradation. This regulator would not be any of the well-known components of the ABA signalling pathway, e.g. the PYR/PYL receptors, PP2C phosphatases or OST1 kinase (Assman and Jegla 2016, Engineer et al., 2016). A future screen for MAX2-interacting proteins using e.g. MAX2 co-immunoprecipitation from isolated guard cells could be used to unravel other components of this specific branch of guard cell signalling and give new information on how different signalling pathways interact to regulate stomatal function.

## Acknowledgements

This study was financially supported by the Academy of Finland Center of Excellence in Molecular Biology of Primary Producers 2014-2019 (grant #307335). C.W. is funded by the Academy of Finland (decision #294580). M.K. received a grant from Svenska kulturfonden (Carin och Gustaf Arppes fond). We acknowledge the support from Doctoral Program in Integrative Life Science (ILS) and Doctoral Programme in Plant Sciences (DPPS). We thank Leena Grönholm for technical assistance in plant growth facilities, Aapo Kalliola for providing a program for automatic leaf area calculation and Pilvi Ackté for technical assistance.

## Supplemental Data

**Supporting Information Figure 1.**
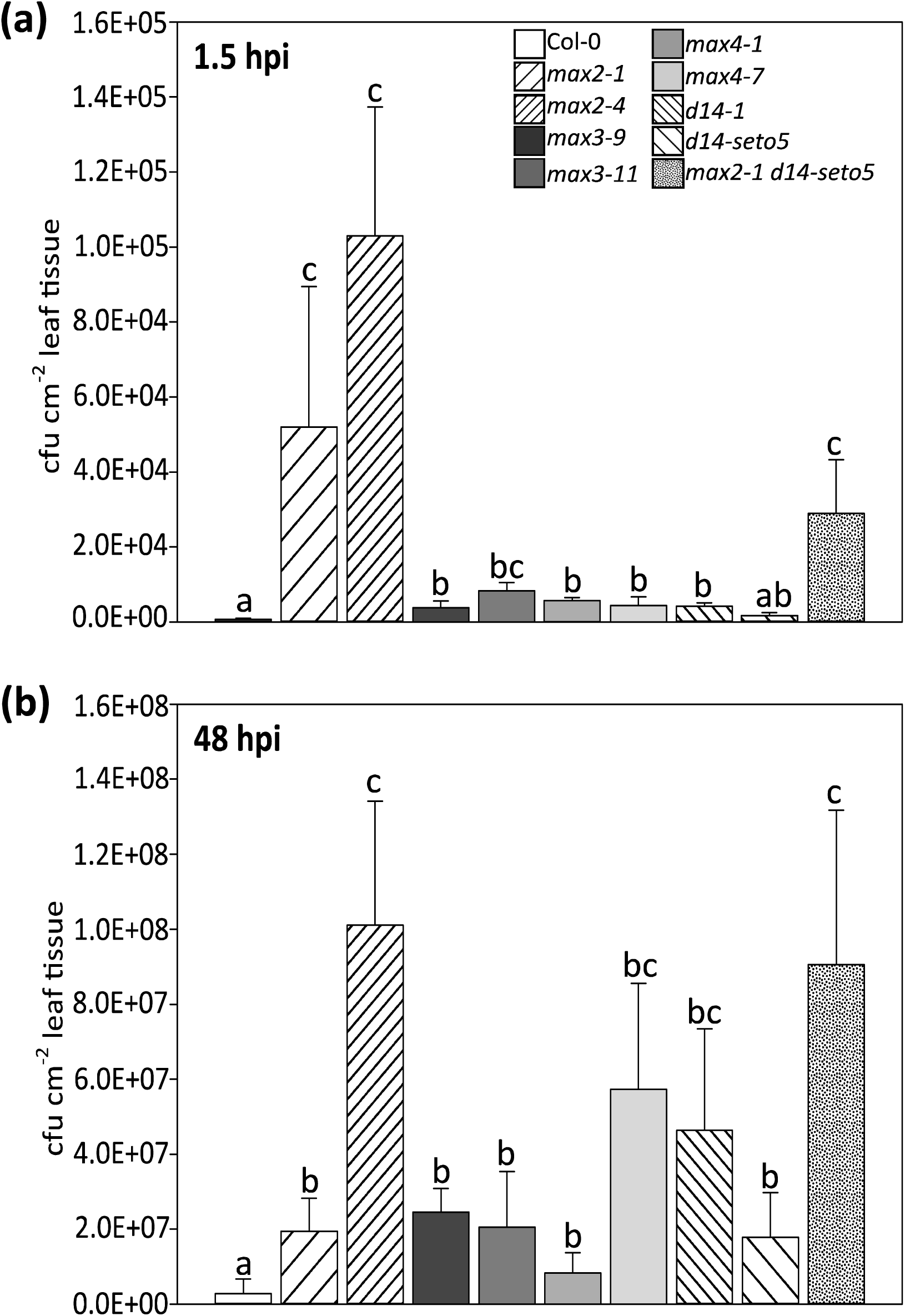
The sensitivity to pathogens in darkness is affected both in strigolactone sensing (*max2*), the receptor (*d14*) and synthesis (*max3* and *4*) mutants. The spray infection with *P. syringae* DC3000 was performed during the dark period of 12 h light rhythm 6 hours after the darkness was applied. In 48 hours post inoculation the infection is in general weaker than when performed in normal light rhythm but otherwise the light-rhythm dependent differences are small. In each experiment four plants per line and three leaves per plant were used to measure the bacterial concentration. The experiment was repeated three times with similar results. The results are shown as means ± SE. In statistical analysis we compared if Col-0 significantly differs from mutants; first a logarithmic transformation was conducted on the data, then univariate analysis of variance combined to Hochberg post hoc test.

**Supporting Information Figure 2.**
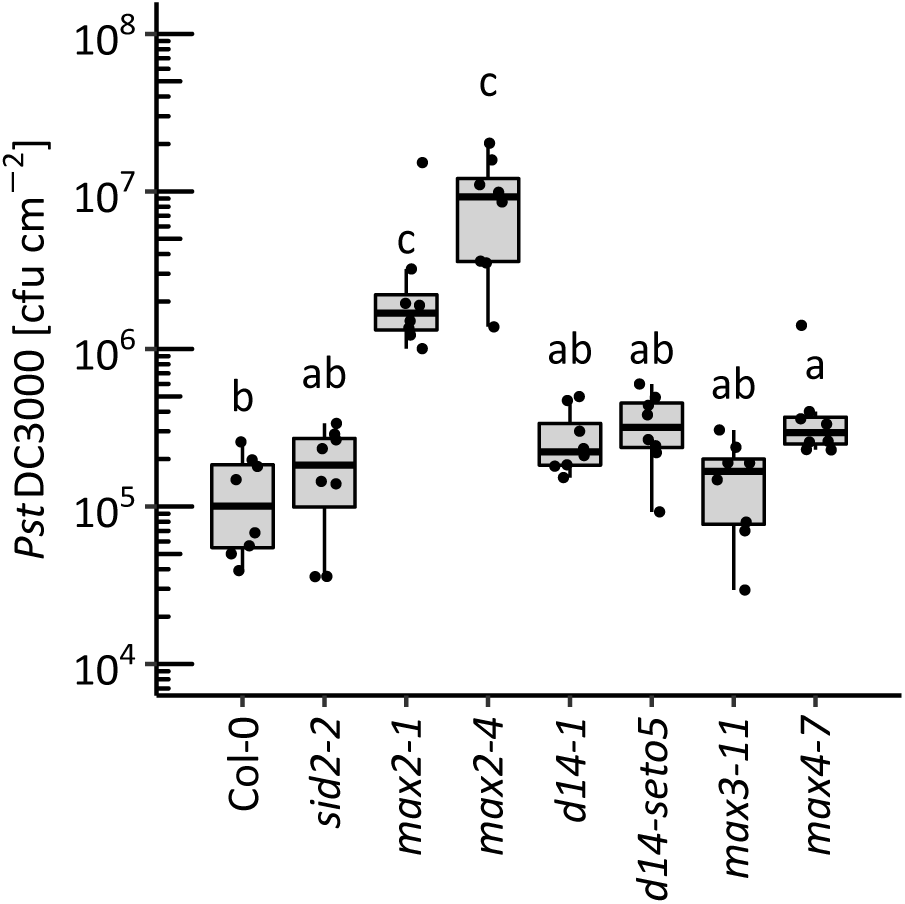
Growth of *Pseudomonas syringae* DC3000 in the same samples used in salicylic acid assay (Figure 2). Bacterial growth was quantified 27 hours after spray infection (OD600 = 0.2) in wild-type (Col-0), strigolactone signalling (*max2-1, max2-4, d14-1* and *d14-seto5*) and biosynthesis mutants (*max3* and *max4*). Salicylic acid biosynthesis mutant (*sid2-2*) was included as a control for no accumulation of free salicylic acid salicylic acid assay. Each dot represents one plant. In total, seven to eight plants were used per line. The experiment were performed three times with similar results. Box plots are summarising data by showing the median, and first and third quartiles. Whiskers are extending to a maximum of 1.5 × interquartile range beyond the box. Different letters indicate significant differences (P < 0.05) as determined by Kruskal-Wallis Rank Sum test followed by Pairwise Wilcoxon Rank Sum tests with multiple testing correction to p-values using Holm method.

**Supporting Information Figure 3.**
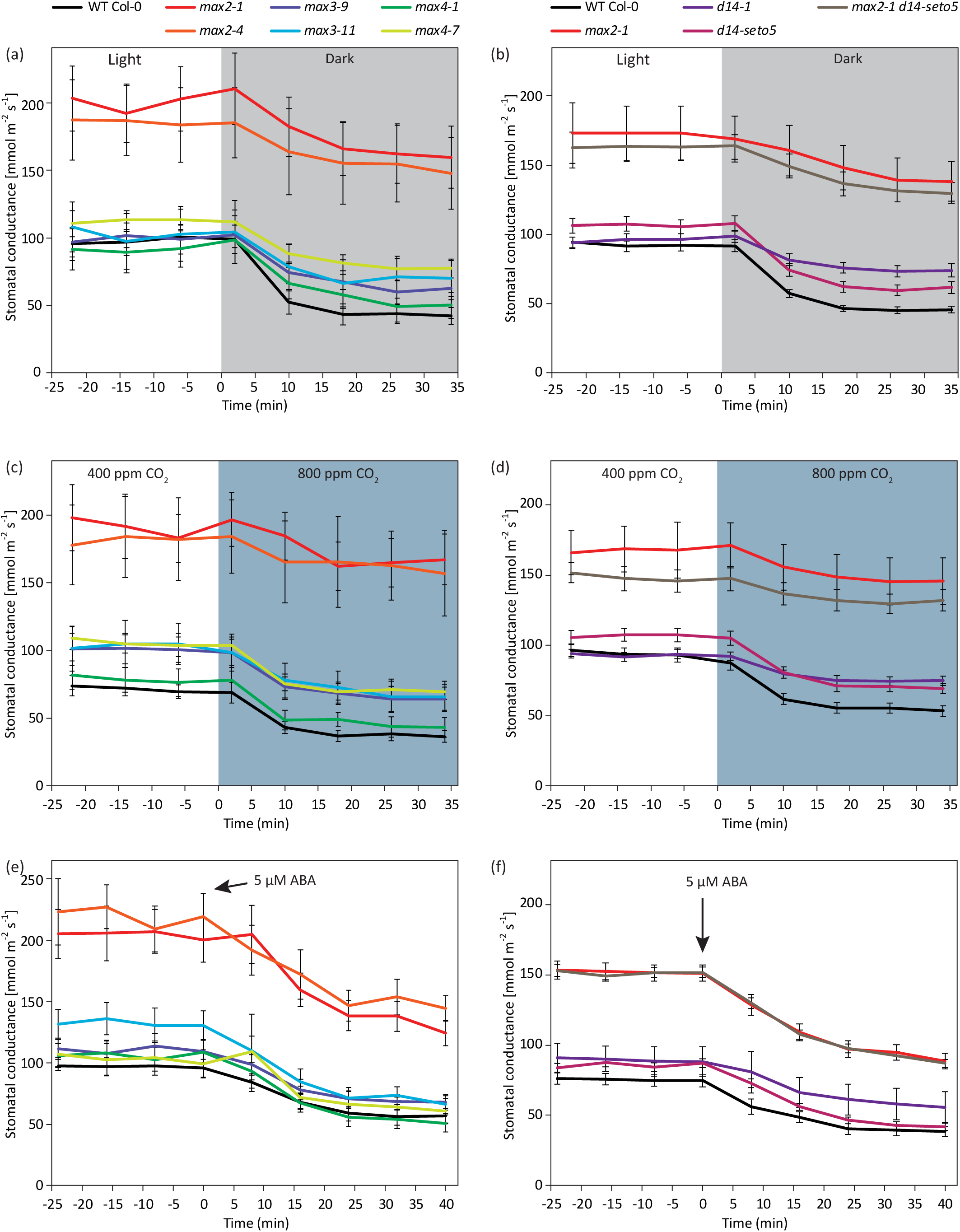
The time course data of stomatal conductance used to calculate the bar graphs of Figure 7c–7h.

**Supporting Information Figure 4.**
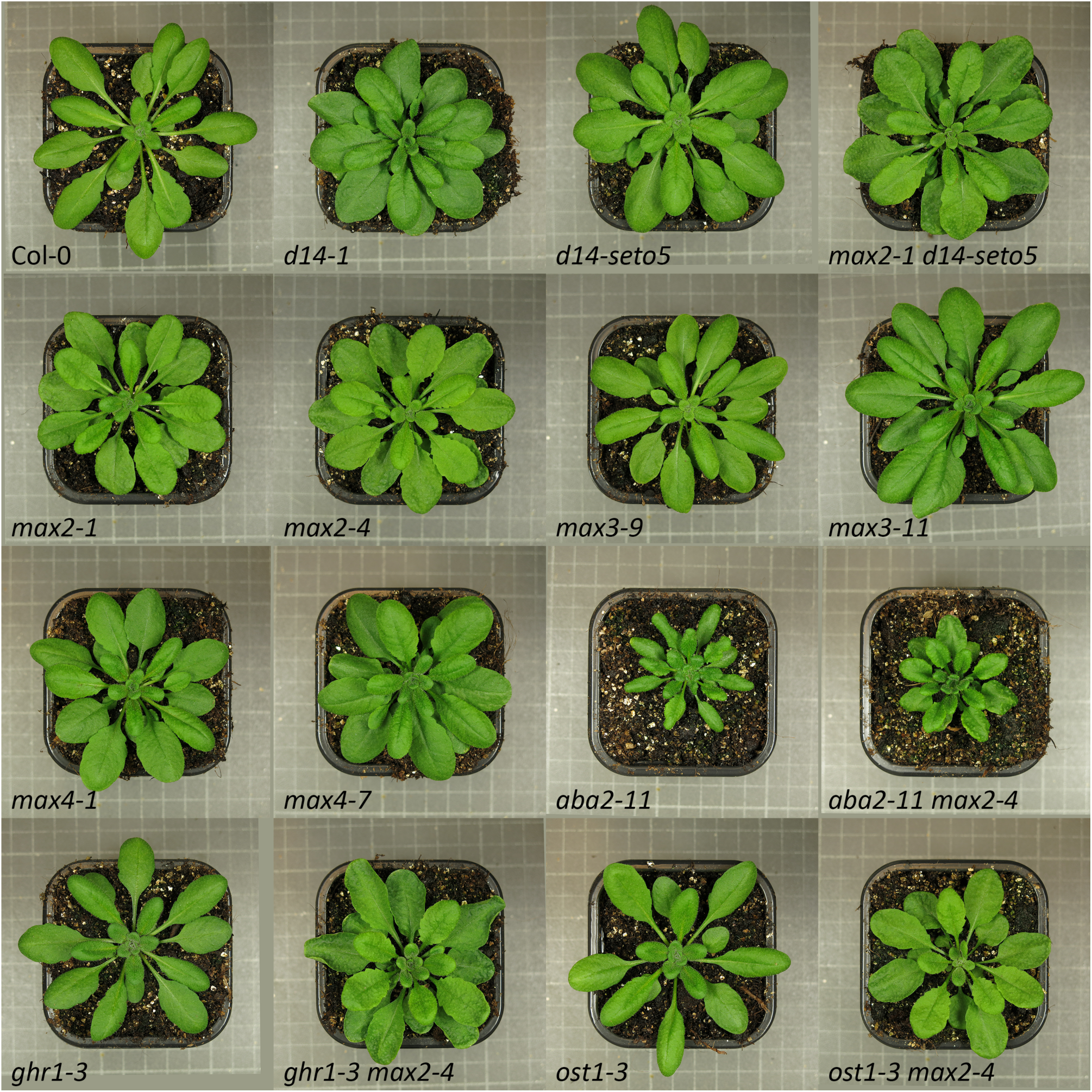
The phenotypes of Arabidopsis Col-0 and the mutant plants involved in the experiments. The phenotype of 4-5 weeks-old plants (Helsinki growth conditions, see Materials and Methods). The size of the growth pot for all the plants is 7×7 cm.

**Supplemental Table 1.**
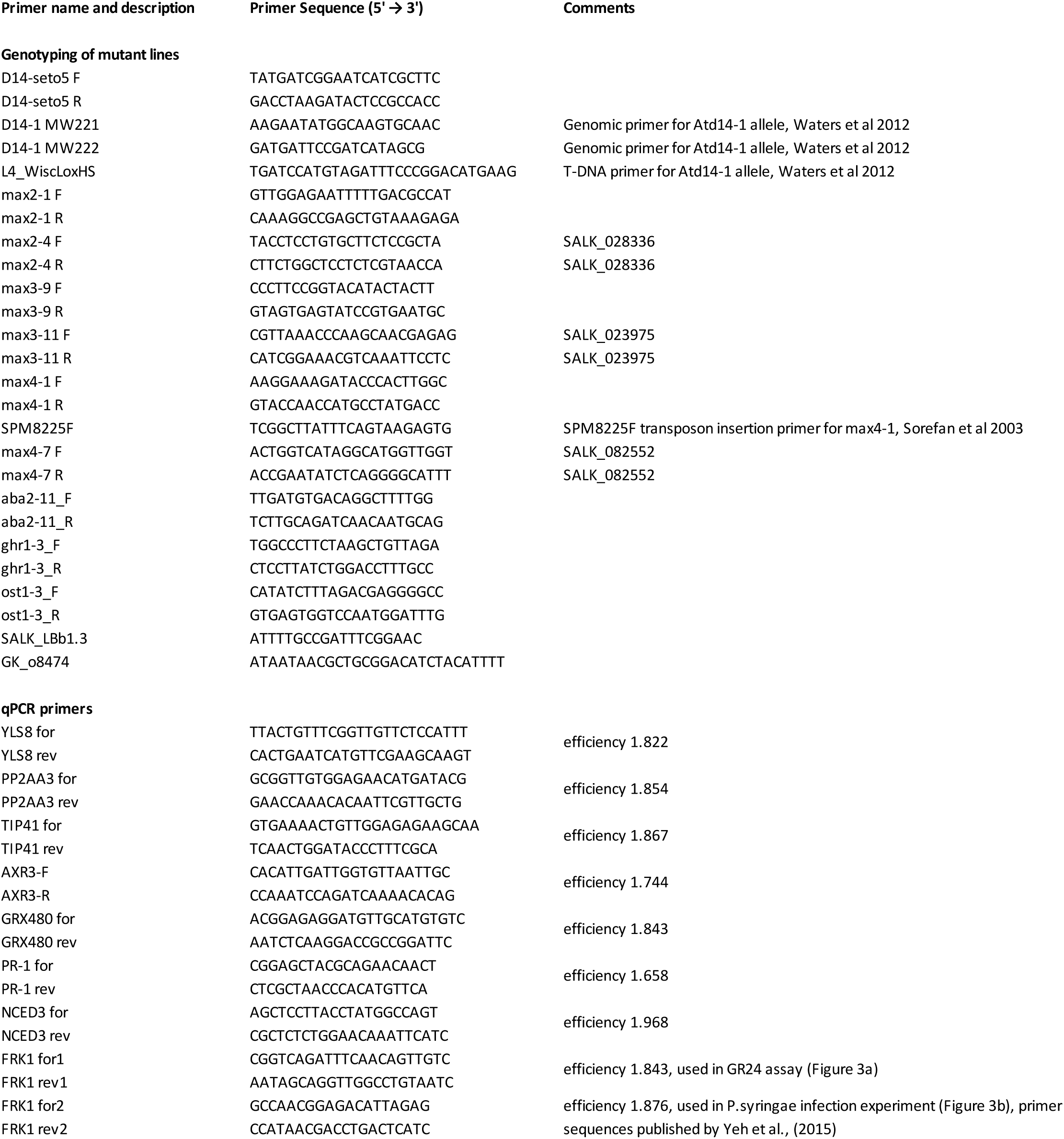
The PCR primers used in genotyping and qPCR primers used in gene expression analysis.

